# Distinctive Features of a Saudi Genome

**DOI:** 10.1101/015909

**Authors:** Ibrahim Alabdulkareem, Anthony J. Robertson, Sami Al Garawi, Mohammed Aljumah, Saeed Al-Turki, Mohammed A. AlBalwi

**Author notes:** **Correspondence to:** Dr. Mohammed A. AlBalwi, Department of Pathology and Laboratory Medicine, MC 1122, King Abdulaziz Medical City, P.O. Box 22490, Riyadh 11426, Kingdom of Saudi Arabia, Facsimile numbers.

## Abstract

We have fully sequenced the genome of an individual from the region of Saudi Arabia. In order to facilitate comparative analysis, an initial characterization of the new genome was undertaken based on single nucleotide polymorphism (SNP). The SNP data having associated population statistics, essentially the HapMap, served to identify features that were rare by comparison. Methods were developed and applied to tag observed SNPs as different and were extended to identify strings or clusters of difference in the individual relative to comparison populations to effectively increase the significance over single SNP comparison. Difference strings identified in the individual relative to each comparison population showed a genome location pattern with various levels of overlap between the comparison populations. The SNP frequencies from the HapMap population samples Ceu and Yri showed a difference inversion relative to the sample genome. The total SNP difference count was greatest between the individual and the Yri population sample while the number and total span of SNP difference clusters was greatest in comparison with the Ceu population sample. The final pattern of difference clusters has served to define distinctive features in the individual genome toward preliminary characterization.

## INTRODUCTION

There is a recognized and promising trend in human health studies involving the use of high-resolution genetic information to compliment diagnostics (Marian 2011; Cooper et al. 2011). High-resolution information based on Microarray (Roberts et al. 2010) and parallel-sequencing (Pareek et al. 2011) methods provide a gain in feature density which may commonly expand the potential of clinical genetics and its impact on health care.

In the process, practical challenges such as handling and analyzing of large data sets are introduced (Pare 2010; Moore et al. 2010). Genome–wide association studies (GWAS) adapt genetic principles to high-resolution feature measurements in the search for disease-associated, not necessarily causal, genetic variations (Kingsley 2011). Comprehensive patient studies have revealed the power of GWAS to define distinctive genetic features, or variations within disease individuals relative to the host population (Visscher et al. 2012).

For a successful outcome to the search for distinctive features, a comparison among a significant number of individuals is required to eliminate personal variations and confidently link any consistent variation to the disease (Gamazon et al. 2012). The human genome is complex and the variation that might have been naively considered as limited in times just after the first human genome release is proving to be more general than expected. The catalogs and collections of variation are becoming extensive (1000 Genomes Project Consortium et al. 2010; Buchanan et al. 2012).

In addition to the challenge of developing a systematic format of variations that naturally increase as more individual genomes are sampled, there is a balancing demand for representing all found variation within a conventional reference as a basis for clinical interpretations.

In what might be a contributing element to an emerging trend in clinical practice, possibly even a paradigm shift (Kuhn 1970); there is an alternate and parallel approach to GWAS. The mapping of rare alleles to disease is centered on the measurements of populations and restricted to high coverage of information rich variations, such as protein coding regions of the genome (Tennessen et al. 2012). A susceptibility allele might have provided some selective advantage in the past but remains, at least with low frequency, due to insufficient generation passage or weak purifying selection. The paradigm of evolutionary medicine and efforts to include high-resolution genetic information demands sufficient sampling and clear understanding of the origins and demographics of rare and disease causing alleles (Nesse et al. 2012).

For the present study, we have reviewed genomic data from a geographic region with great opportunities for the application of medical genetics as well as population studies (Tadmouri et at. 2006; Armitage et al. 2011). An individual, healthy Saudi Arabian male was the contributor of DNA sequenced to moderate coverage as well as being analyzed for SNPs with a Microarray method. The amount of sequence data has provided the starting point for in depth mapping of variation with additional full-genome data expected to become available through 1000 genomes (Hammond 2008).

For the present, we have mapped the SNPs and placed the found haplotype into the context of available populations with significant measure of allele overlap and frequency to draw upon for statistical analysis. Particular attention was focused on genetic difference that is unique to the sample and distinguished the sample within the comparison.

## METHODS

### Clinical

The purified genomic DNA collected from a peripheral blood of a healthy male individual was used as a source material for genomic analysis, both Sequencing and Microarrays. Paired-read sequence data includes a total of 43 flow-cell lanes which were produced from 3 separate genome fragment libraries following standard protocols for the Illumina GAx (Mate Pair Library Sample Preparation Guide). DNA from the same source was used to develop Affymetrix SNP chips 6.0 following the vendor-supplied protocols for target labeling and scanning (Human SNP assay 6.0 User Guide for Automated Target Preparation Affymetrix).

### Sequence Assembly

Genome version GRCh37/hg19 was downloaded onto a standard workstation by HTTP directly from NCBI Entrez (NCBI website). Download options were selected for each chromosome, including the mitochondrial to contain the whole sequence in forward orientation with default annotation features in GenBank-full format. The reference sequences were subsequently imported without changes into the assembly software, Genomics Workbench (Genomics Workbench User Manual).

The sequence read data were imported into the Genomics Workbench as 43 separate files with undetermined fragment length (mate-pair import) so that alignment parameters could be optimized independently for each data set. Prior to assembly, the reads were trimmed based on quality scores using the Workbench Trim Tool to allow for global alignment that extends the mismatch penalty to the ends of reads during alignment (Genomics Workbench User Manual). An average of 1.02% of the total read length was trimmed from the data that ranged in length from 39 to 75 as determined by the instrument cut-off value used. A data sample for each of the 43 files was pre-assembled ignoring non-specific reads to provide a paired read length distribution that was in-turn used to set the optimal alignment parameters.

The alignment to reference was performed using default settings for both long and short reads (short<56, others are long) with the Global Alignment option and paired parameters set to range from the average (<u>+/-</u>) 2.5 standard deviations based on the predetermined pair fragment length.

### Variant Mapping

SNP detection was performed on the complete assembly using the SNP detection tool available in the Genomics Workbench. The settings for quality were Window length = 11, Maximum gap and mismatch count = 2, Minimum central quality = 20 and a Minimum average quality = 15. The settings for significance were Minimum coverage = 3 and Minimum variant frequency (%) = 35.0. The Maximum expected variations (ploidy) was set as 2. The output of Genome Workbench SNP detection is a table of the SNPs detected in the aligned data relative to the reference and that pass the settings filter for quality and significance.

Affymetrix SNP chip 6.0 measurements from the same sample were compared to the SNP mapping results obtained through sequencing, assembly and SNP detection. For this comparison, it is the positional information relative to the reference sequence that is used to register the non-identical results files. Overlapping, common positions were determined with a Venn diagram based online tool (Oliveros 2007). Each separate position list was compensated for missing positions present in the comparison list leading to the generation of a single, matched comparison table. A simple positive scoring that required absolute agreement between sequence and hybridization based methods was used after all positions without full adjacent information were removed through applied filters. No corrections were made for major and minor alleles or the level of confidence of the SNP call so that positive scores were assigned to high-confidence matches only.

### Population Frequency Tables

The distinction value of the SNPs identified in the reference assembly was extended through comparison to population frequencies. The variant annotation files available to download from the UCSC Table Browser (UCSC genome Bioinformatics Group; Karolchik et al. 2004; Fujita et al. 2011) in .gff format with a corresponding frequency table representing a population, essentially the HapMap SNPs, were added to the reference in the Genomics Workbench as separate annotation tracks with an available software plug-in designed for that purpose (Annotate with GFF file User Manual CLC bio). More relevantly, the tables of SNP population frequencies for each of the populations Ceu, Chb, Jpt and Yri were combined and filtered for complete overlap with the found SNPs to produce a single frequency table for each chromosome representing the individual SNP and each population. An online Venn diagram tool was used to find the intersection among the combined data sets based on the consistent numbering relative to the reference genome (Oliveros 2007). The four selected populations provide substantial sampling of SNPs and individuals that decreases significantly if additional population tables are included.

### Determine Distinctive Variations

The distinctive features of the individual genome based on SNP variation were identified by two separate methods. Custom “R” scripts directed a global approach to find those SNPs in the frequency tables that are either frequent or rare in a single population only. An allele was considered to be rare in a population if the frequency was less than 10% and frequent if 50% or greater. The values used in allele calling were empirically determined to return a workable sampling.

Measure of difference of a population relative to the individual (or other populations) was counted as the frequent or rare SNP unique to the population after normalizing to the total number of frequent or rare alleles within that population. A directed approach was also taken to identify distinctive SNPs by making a set of rules to call the found SNP different from the corresponding one in each population (Supplemental Fig. 1). The measure of difference was directly counted as the total number of difference calls.

### Determine Distinctive Regions

The identification of difference strings was again performed in the frequency table described above with the addition of the difference tags that were added. For the purpose of identification of strings of sequential difference, the frequency table was ordered from lowest to highest position. A script was developed to report the start and end positions of the string as well as the number of SNPs within the string. The script was run to identify all strings and filtered to strings of at least 10 SNPs relative to each of the comparison populations. An annotation file in .gff format was prepared for the difference strings identified in each chromosome with a separate annotation types given to each comparison population. Annotation files generated were added to the reference genome in the Genome Workbench. Unique features were searched by manually stepping through the graphics pane of the Genomics Workbench and viewing the overlap within the difference annotation, a process aided with the search for annotation tools available in the Workbench.

## RESULTS

### Assembly of the sequence data to the reference

The total length of the current human male reference genome is 3,112 Mb with 2,882 Mb outside of regions annotated as “Gap”. For this study an individual Saudi male genome sample was sequenced resulting in 42,996 Mb of paired-read sequence as data source in an assembly to the reference. With the assembly parameters that were used most of the reads, 97%, were included in the final assembly for 14.5 fold sequence coverage of the non-gap reference (Table 1). The 3% of the reads that did not map during the assembly process were saved and de-novo assembly was performed using the de-novo tool in the assembly software. About 7% of the unmapped reads were assembled into contigs of length at least 2.5 times the average read length as a result of de-novo assembly. Only 2 of the contigs produced from de-novo assembly have a non-human top blastN hit to NCBI-nt and those 2 were both more similar to bac clones from *Pan troglodytes* relative to other GenBank entries. An average across all chromosomes of the reads that mapped to the reference as paired reads was 92% ranging from a low of 73% on the X chromosome to a high of 96% for chromosome 12 and also for the Mitochondrion reference. Considering the entire fragment length or paired read distance, the physical coverage of the reference was 28.90 (Table 1).

**Table 1:**
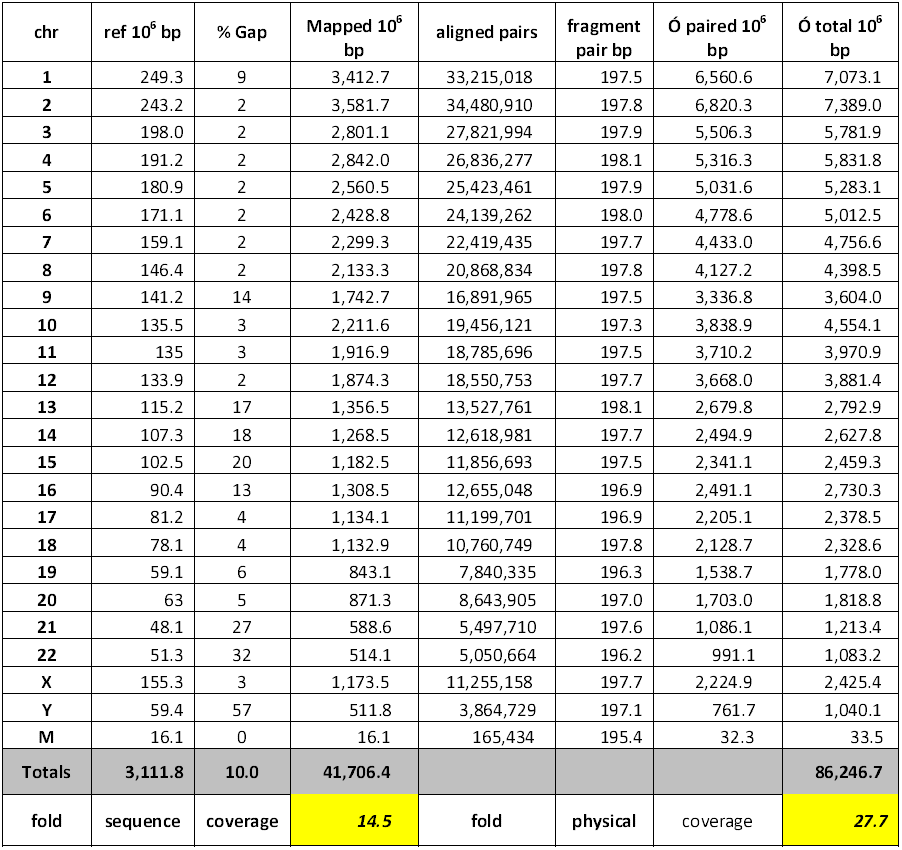
Mapping Statistics

| chr | ref 10<sup>6</sup> bp | % Gap | Mapped 10<sup>6</sup> bp | aligned pairs | fragment pair bp | Ó paired 10<sup>6</sup> bp | Ó total 10<sup>6</sup> bp |
| --- | --- | --- | --- | --- | --- | --- | --- |
| 1 | 249.3 | 9 | 3,412.7 | 33,215,018 | 197.5 | 6,560.6 | 7,073.1 |
| 2 | 243.2 | 2 | 3,581.7 | 34,480,910 | 197.8 | 6,820.3 | 7,389.0 |
| 3 | 198.0 | 2 | 2,801.1 | 27,821,994 | 197.9 | 5,506.3 | 5,781.9 |
| 4 | 191.2 | 2 | 2,842.0 | 26,836,277 | 198.1 | 5,316.3 | 5,831.8 |
| 5 | 180.9 | 2 | 2,560.5 | 25,423,461 | 197.9 | 5,031.6 | 5,283.1 |
| 6 | 171.1 | 2 | 2,428.8 | 24,139,262 | 198.0 | 4,778.6 | 5,012.5 |
| 7 | 159.1 | 2 | 2,299.3 | 22,419,435 | 197.7 | 4,433.0 | 4,756.6 |
| 8 | 146.4 | 2 | 2,133.3 | 20,868,834 | 197.8 | 4,127.2 | 4,398.5 |
| 9 | 141.2 | 14 | 1,742.7 | 16,891,965 | 197.5 | 3,336.8 | 3,604.0 |
| 10 | 135.5 | 3 | 2,211.6 | 19,456,121 | 197.3 | 3,838.9 | 4,554.1 |
| 11 | 135 | 3 | 1,916.9 | 18,785,696 | 197.5 | 3,710.2 | 3,970.9 |
| 12 | 133.9 | 2 | 1,874.3 | 18,550,753 | 197.7 | 3,668.0 | 3,881.4 |
| 13 | 115.2 | 17 | 1,356.5 | 13,527,761 | 198.1 | 2,679.8 | 2,792.9 |
| 14 | 107.3 | 18 | 1,268.5 | 12,618,981 | 197.7 | 2,494.9 | 2,627.8 |
| 15 | 102.5 | 20 | 1,182.5 | 11,856,693 | 197.5 | 2,341.1 | 2,459.3 |
| 16 | 90.4 | 13 | 1,308.5 | 12,655,048 | 196.9 | 2,491.1 | 2,730.3 |
| 17 | 81.2 | 4 | 1,134.1 | 11,199,701 | 196.9 | 2,205.1 | 2,378.5 |
| 18 | 78.1 | 4 | 1,132.9 | 10,760,749 | 197.8 | 2,128.7 | 2,328.6 |
| 19 | 59.1 | 6 | 843.1 | 7,840,335 | 196.3 | 1,538.7 | 1,778.0 |
| 20 | 63 | 5 | 871.3 | 8,643,905 | 197.0 | 1,703.0 | 1,818.8 |
| 21 | 48.1 | 27 | 588.6 | 5,497,710 | 197.6 | 1,086.1 | 1,213.4 |
| 22 | 51.3 | 32 | 514.1 | 5,050,664 | 196.2 | 991.1 | 1,083.2 |
| X | 155.3 | 3 | 1,173.5 | 11,255,158 | 197.7 | 2,224.9 | 2,425.4 |
| Y | 59.4 | 57 | 511.8 | 3,864,729 | 197.1 | 761.7 | 1,040.1 |
| M | 16.1 | 0 | 16.1 | 165,434 | 195.4 | 32.3 | 33.5 |
| **Totals** | **3,111.8** | **10.0** | **41,706.4** |  |  |  | **86,246.7** |
| **fold** | **sequence** | **coverage** | **14.5** | **fold** | **physical** | **coverage** | **27.7** |

### SNP mapping and accuracy

With high confidence in the outcome of the sequencing and assembly process; the focus was naturally directed to the variation that might be revealed in the assembly, an interest due to the under-sampled geographic region of the sample genome. Even with all of the possible types of variation and having available tools to map different types of variation, deliberate attention was directed to SNPs. The advanced organization and genome coverage of the HapMap and the availability of SNP frequency tables within representative populations were considered as an information-rich source for a comparison between the sample genome and external population alleles. The lack of quantitative information for much of the reported variation supported the attention given to HapMap SNP data. A total of 3,291,451 SNPs were mapped in the assembly with the tool and parameters described (methods). Accuracy of SNP mapping was assessed by comparison between the SNP mapping and the results from Affymetrix SNP chip 6.0 developed from the same sample. The overlap between the found SNPs and Affymetrix hybridization probes was 10.6% and those SNPs were used to test the platform agreement. The average agreement determined was 92.7% with a range from 90.4% to 99.1% across the reference (supplemental Table 1).

### Comparison of found SNPs to HapMap populations

A frequency table was built after selecting the intersection of the HapMap SNPs that are present as called SNPs in the alignment of the individual to the reference. The side-by-side genome alignment was thus extended into a statistical variation space. The SNP sample size found in the assembly and filtered for presence in all included HapMap populations is 1.4 times greater than the total Affymetrix chip SNP features. The average population sample size for each SNP is 93 to provide significant sample dimensions (supplemental Table 2). Including additional HapMap populations in the frequency tables reduced the sample size due to fewer SNPs tested in those studies. Only the 4 populations with a large, overlapping SNP test pool and with large genome sampling sizes were considered to provide an optimal sampling dimension.

Difference measures were applied to contrast the sample genome against HapMap populations representative of geographic or ethnic groups (Barnes 2006). Search results for rare or frequent alleles present within a single population were used to define unique representation of the allele in a single population relative to other populations. This rare or frequent difference measure is a 2 *vs*. 1 approach since the Chb and Jpt population statistics were excluded due to an introduced similarity bias. The data were normalized to the total number of frequent and rare alleles to again avoid bias, in this case the potential bias caused by having a higher number of such rare or frequent alleles within their population. A consistent pattern of the sample genome having a greater difference to the Yri population was observed following this measure (Fig. 1). An alternate approach was developed to produce a decision of difference for each sample SNP. A formula was developed and applied to each SNP to compare the sample against each population. The formula produces a different-or-not decision relative to corresponding population frequency observed among possible alleles. In many cases the decision is as clear as a homozygous allele in the sample not found in the comparison population and in some cases the decision is marginal. The optimized formula and large sample size do provide a detail level useful for distinction among similar populations. The consistent pattern observed with a 2 *vs*. 1 approach of counting the presence of rare or frequent alleles in a single population is also found with the decision formula method. The Yri population contained more differences from the sample genome and the Ceu population had the fewest differences relative to the found SNPs (Fig. 2).

**Figure 1:**
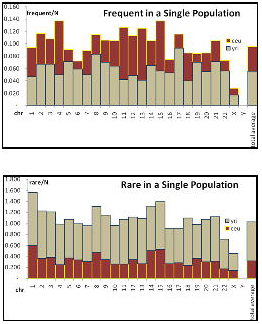
A summary of the total difference count based on the single population method. The top panel shows the

**Figure 2:**
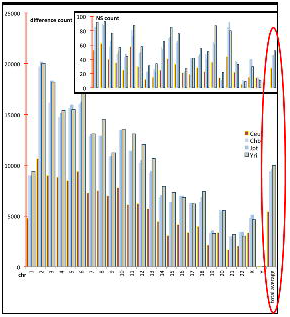
A summary of the total difference count based on a formula method. The formula applied to each SNP tested provides a difference decision. The results of the difference call formula are shown as the total number of differences for each population separated by chromosome. The inset in the graph is the subset of those SNPs that are different and are also non-synonymous.

### Difference strings distinguish unique genomic regions

In the process of labeling the relative difference of each SNP present in the frequency tables uninterrupted strings of difference labels were observed. Based on this observation of strings of difference labels in consecutive positions, these were investigated further as possible chromosomal regions of concentrated differences. We retrieved all such strings of at least 10 consecutive difference-labeled SNPs. The difference strings were in turn added as annotation tracks to the reference genome. Difference strings relative to each comparison population were kept as separate annotation types. The population samples Ceu and Jpt had the largest number of strings with a corresponding large and mainly overlapping span (Fig. 3). A difference inversion was observed concerning the strings present in the Yri population. The span of strings was found to be less in the Yri population than in other populations contrary to the observation that the total number of differences was greatest in that population. The average string length over all chromosomes was 15.2 consecutive SNPs.

The regions of the genome where the annotation of difference strings overlapped were visually identified in the Genomics Workbench graphics display following the addition of annotations representing the difference strings from each comparison population. There were 46 separate regions across the genome having overlapping difference string annotation (Table 2). The regions with such overlapping annotation are unique and distinguish the sample genome as different from all populations in these regions for the SNP set used.

**Figure 3:**
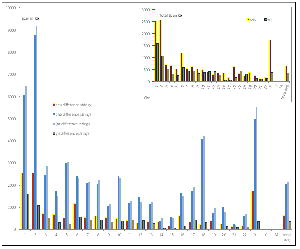
A summary of the total span of difference strings. Consecutive SNPs that are different from the comparison population and forming a string of length 10 or greater are represented. The span of a represented string is the distance between the end positions. The total span is scaled in the inset for 2 populations only.

**Figure 4:**
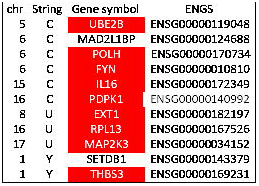
Genes containing at least 3 overlapping difference strings that are associated in an interaction network as defined through bioGRID and identified using g:Profiler with an assigned significance of p=0.00206. The associated genes were identified pairwise by adding the list of unique genes listed in Table 3 to the list of either Yri-Chb-Jpt (Y) or Ceu-Chb-Jpt (C). The Y list and the C list of genes did not have any significant functional linkage assigned using g:Profiler with default settings for significance. Combining the lists extended the bioGRID network by the 2 genes indicated.

**Table 2:** Genes that contain overlapping difference strings

| unique |  | Yri, Chb & Jpt |  |  |  | CeU, Chb & Jpt |  |  |  |
| --- | --- | --- | --- | --- | --- | --- | --- | --- | --- |
| chr | gene | chr | gene | chr | gene | chr | gene | chr | gene |
| 3 | ABI3BP | 1 | ST3GAL3 | 2 | WDPCP | 2 | CAD | 7 | ELMO1 |
| 4 | SCOC | 1 | DMBX1 | 3 | ADH4 | 2 | SLC30A3 | 7 | CLEC2L |
| 4 | CLGN | 1 | AKNAD1 | 3 | LOC100507053 | 2 | DNAJC5G | 9 | PTPRD |
| 5 | GUSBP1 | 1 | SETDB1 | 3 | LSM6 | 2 | TRIM54 | 10 | KIAA1217 |
| 6 | PRIM2 | 1 | MUC1 | 4 | FAM81B | 2 | MPV17 | 10 | PAX2 |
| 7 | CACNA2D1 | 1 | THBS3 | 5 | TULP4 | 2 | LRRTM4 | 11 | RCN1 |
| 8 | ZFPM2 | 1 | MTX1 | 6 | ITGB8 | 2 | THSD7B | 11 | PTPRJ |
| 8 | EXT1 | 1 | GBA | 6 | CHN2 | 2 | GALNT13 | 11 | FAT3 |
| 9 | PTPRD | 1 | FAM189B | 6 | NUDCD3 | 2 | CCDC148 | 12 | ADAMTS20 |
| 10 | TDRD1 | 1 | SCAMP3 | 6 | COBL | 2 | PARD3B | 12 | PTPRQ |
| 13 | ZMYM5 | 1 | CLK2 | 6 | BRI3 | 2 | UNC80 | 13 | CDK8 |
| 16 | HYDIN | 1 | HCN3 | 6 | BAIAP2L1 | 3 | ERC2 | 13 | TMCO3 |
| 16 | SPG7 | 1 | PKLR | 6 | CHRM2 | 3 | GXYLT2 | 15 | GANC |
| 16 | RPL13 | 1 | FDPS | 6 | CNTNAP2 | 3 | SLC41A3 | 15 | PTPLAD1 |
| 17 | MAP2K3 | 1 | RUSC1 | 7 | PNOC | 3 | KCNAB1 | 15 | C15orf44 |
| 22 | CCDC157 | 1 | ASH1L | 7 | LACTB2 | 3 | PIK3CA | 15 | IL16 |
|  |  | 1 | OR10K1 | 7 | XKR9 | 4 | SORCS2 | 15 | NTRK3 |
|  |  | 1 | PRRX1 | 8 | ZNF248 | 4 | LOC100505875 | 16 | TBC1D24 |
|  |  | 1 | KCNT2 | 8 | LIPK | 4 | TNIP3 | 16 | AMDHD2 |
|  |  | 1 | TRAF3IP3 | 12 | PDS5B | 5 | DHFR | 16 | CEMP1 |
|  |  | 1 | IRF6 | 13 | SLC28A2 | 5 | MTRNR2L2 | 16 | PDPK1 |
|  |  | 1 | USH2A | 15 | SPECC1 | 5 | MSH3 | 16 | PKHB |
|  |  |  |  | 16 | MAPT | 5 | SKP1 | 16 | LOC100507534 |
|  |  |  |  | 16 | KIAA1267 | 5 | CDKL3 | 16 | HYDIN |
|  |  |  |  | 16 | LOC644669 | 5 | UBE2B | 17 | OR1G1 |
|  |  |  |  | 19 | SEC14L4 | 5 | BTNL3 | 17 | ADAP2 |
|  |  |  |  |  |  | 6 | XPO5 | 17 | AATF |
|  |  |  |  |  |  | 6 | POLH | 17 | FLJ37644 |
|  |  |  |  |  |  | 6 | GTPBP2 | 19 | LOC100507550 |
|  |  |  |  |  |  | 6 | MAD2L1BP | 19 | GMFG |
|  |  |  |  |  |  | 6 | RSPH9 | 19 | SAMD4B |
|  |  |  |  |  |  | 6 | C6orf142 | 19 | PSG3 |
|  |  |  |  |  |  | 6 | MRAP2 | 19 | PSG8 |
|  |  |  |  |  |  | 6 | KIAA1009 | 22 | SGSM1 |
|  |  |  |  |  |  | 6 | LACE1 | 22 | CACNG2 |
|  |  |  |  |  |  | 6 | FYN | 22 | TNRC6B |
|  |  |  |  |  |  |  |  | X | TRPC5 |
|  |  |  |  |  |  |  |  | X | ZCCHC16 |
|  |  |  |  |  |  |  |  | X | HTR2C |
|  |  |  |  |  |  |  |  | X | IL13RA2 |
|  |  |  |  |  |  |  |  | X | ODZ1 |

Additional tests were performed to further assess the unique regions identified, for chromosomes 17 and 22. Firstly, variation types such as deletions and insertions were found using the available tools in the Genome Workbench. There was no co-occurrence of additional variation types with unique regions identified based on SNP difference strings. Also, the alleles present in the alignment positions of HapMap SNPs not identified as SNPs due to agreement with the reference were recovered. These non-SNPs were added to the original frequency tables and the difference formula was repeated to label these positions in addition to those that were previously labeled. Some changes in the difference string positions resulted from inclusion of non-SNP positions but the changes were balanced and mainly supported the difference strings as before. Slight shifts in the start and end position, interrupted strings that were replaced by new adjacent strings were observed. Finally, found SNPs that were not in HapMap positions were retrieved and searched for clusters. Clusters of found non-HapMap SNPs having near proximity of at least 10/Kb were recovered into .gtf files and added as an additional annotation track. Interestingly, there was very little co-occurrence of non-HapMap, found SNP clusters with HapMap strings. All together these additional tests supported the utility of overlapping HapMap difference strings to successfully identify unique regions where other approaches may have fallen short.

Of the 46 genomic regions that might be considered as unique or distinctive based on overlapping difference string annotation 30 were located within regions annotated as genes. Since some of the genes that contain unique regions contain multiple, separate unique regions (Table 2) a total of 16 genes are affected. Amazingly, the total span of unique regions is 0.016% of the non-gap genome suggesting selective force directing the location of the unique regions to overlap gene regions. The g:Profiler (Reimand et al. 2007; Reimand et al. 2011) was employed to test for functional relations among the genes containing unique regions listed in Table 2. Of the 16 genes listed as having at least one unique region 3 genes are associated with a bioGRID network (Stark et al. 2006; Chatr-Aryamontri et al. 2013) with a g:Profiler significance of 0.00206 using all default settings for significance. Addition of genes with difference strings from at least 3 populations, also listed in (Table 2), to the unique genes expanded the bioGRID when lists were added pairwise. The genes associated in a bioGRID are shown in (Fig. 2).

## DISCUSSION

Finding variation in an alignment of sequence data to a reference genome is meaningless unless the comparison is extended to a larger basis. In the alignment process the reference may become as an introductory step in the comparison of the sample genome to any and all, to uncover the unique qualities of the individual. Steps that follow the assembly process or any other process of initial genome characterization ultimately require a body of comparison features. This well known challenge of forming an adequate basis of comparison remains a true challenge requiring deep organization. The HapMap project has quickly become a classic attempt to the formation of a comparison scale and has seeded a growth in complementary projects that aim to catalog and count variable genomic features. Forming the connections between high-resolution haplotype and health risks remains elusive.

The review of data that was generated from a region with advanced disease tracking methods and a developing interest to adopt high resolution genome analysis in clinical practice has presented an exceptional opportunity to advance the process of connecting the dots, forming the connections between haplotype and health.

We have performed an initial characterization of the variation present in an individual Saudi. With a focus on the subset of found SNPs that are also represented in each of the HapMap representative populations (Ceu, Chb, Jpt and Yri) a basis of comparison was formed. The similarity of the individual to the Ceu population sample as well as the difference between the individual and the Yri population sample based on the total count of SNPs that differed in the comparison took on an inverse when regions of the genome were considered. Strings of SNPs that were adjacent to each other and different from the comparison population were more numerous and covered more of the genome in the Ceu sample than in the Yri population. The co-occurrence or overlap of difference strings marked unique features in the individual and focused attention to a relative limited space within a vast sea of sequence.

It is noteworthy that the SNPs in difference strings were closer together on the average relative to the overall density of SNPs (Table 3). The average SNP density is consistently higher in the span of identified difference strings relative to the average SNP density across the entire genome. The string density is higher across all chromosomes and for difference strings relative to each of the comparison populations. For the majority the average difference string density is even higher than the density of the total SNPs found before filtering to HapMap overlap. With this high density established, the difference strings become concentrated regions of difference. This non-random distribution has been noted before (Amos 2010) and various explanations have been explored. The natural force or biological process that leads to the accumulation of SNPs that are rare or different by comparison into clusters or strings would also undoubtedly generate rare or different phenotypic character.

**Table 3:** SNP density

| chr | non-gab Kb | total found | total/Kb | actual sample<br>(HM_overlap) | sampled/Kb | ceu per Kb in<br>clusters | chb per Kb<br>in clusters | jpt per Kb<br>in clusters | yri per Kb in<br>clusters |
| --- | --- | --- | --- | --- | --- | --- | --- | --- | --- |
| 1 | 226,281 | 378,672 | 1.67 | 102,926 | 0.455 | 0.6793 | 0.7116 | 0.6593 | 0.6956 |
| 2 | 238,357 | 419,728 | 1.76 | 106,184 | 0.445 | 0.9855 | 0.8036 | 0.8052 | 1.3648 |
| 3 | 194,797 | 198,278 | 1.02 | 91,796 | 0.471 | 1.5042 | 1.0451 | 1.1055 | 1.4785 |
| 4 | 187,712 | 219,532 | 1.17 | 84,230 | 0.449 | 1.6035 | 1.452 | 1.7539 | 2.2272 |
| 5 | 177,745 | 180,454 | 1.02 | 79,603 | 0.448 | 2.6888 | 1.5062 | 1.5775 | 3.4779 |
| 6 | 167,745 | 186,211 | 1.11 | 82,165 | 0.490 | 1.6799 | 1.3185 | 1.2816 | 1.9098 |
| 7 | 155,854 | 164,975 | 1.06 | 66,878 | 0.429 | 1.4369 | 1.0832 | 1.1950 | 1.5258 |
| 8 | 142,939 | 157,322 | 1.10 | 71,300 | 0.499 | 1.3652 | 1.2574 | 1.3570 | 2.1824 |
| 9 | 121,043 | 123,218 | 1.02 | 55,856 | 0.461 | 1.7944 | 1.3873 | 1.4009 | 2.0038 |
| 10 | 131,715 | 151,480 | 1.15 | 65,289 | 0.496 | 2.2937 | 1.2404 | 1.4740 | 2.2010 |
| 11 | 131,330 | 147,017 | 1.12 | 63,285 | 0.186 | 1.4145 | 1.391 | 1.5809 | 1.5865 |
| 12 | 130,532 | 134,261 | 1.03 | 56,993 | 0.437 | 1.7270 | 1.2142 | 1.2023 | 1.4230 |
| 13 | 95,590 | 111,491 | 1.17 | 51,686 | 0.541 | 1.1086 | 1.1026 | 1.2691 | 1.7398 |
| 14 | 88,290 | 92,810 | 1.05 | 41,135 | 0.466 | 1.3538 | 1.275 | 1.4556 | 1.6482 |
| 15 | 82,221 | 82,667 | 1.01 | 36,887 | 0.449 | 1.4823 | 1.5122 | 1.0930 | 2.8237 |
| 16 | 78,935 | 94,347 | 1.2 | 35,910 | 0.455 | 0.9273 | 1.4018 | 1.0939 | 1.9512 |
| 17 | 77,895 | 75,332 | 0.97 | 28,901 | 0.371 | 0.8620 | 1.1454 | 1.1216 | 1.5868 |
| 18 | 74,757 | 86,779 | 1.16 | 39,313 | 0.526 | 0.9051 | 1.2038 | 1.0757 | 1.3467 |
| 19 | 55,859 | 60,106 | 1.08 | 17,805 | 0.319 | 0.9418 | 0.7158 | 0.6043 | 3.9231 |
| 20 | 59,606 | 63,417 | 1.06 | 27,490 | 0.461 | 1.1664 | 1.3087 | 1.2722 | 1.1194 |
| 21 | 35,159 | 46,564 | 1.32 | 16,774 | 0.477 | 1.7390 | 1.5516 | 1.4587 | 1.6553 |
| 22 | 34,895 | 38,003 | 1.09 | 14,489 | 0.415 | 1.4832 | 1.3504 | 1.1733 | 1.1536 |
| X | 151,351 | 64,308 | 0.42 | 24,457 | 0.162 | 0.6066 | 0.6587 | 0.5814 | 1.3335 |
| Y | 25,754 | 13,690 | 0.53 | 69 | 0.003 | 0.0000 | 0.0000 | 0.0000 | 0.0000 |
| M | 16,097 | 31 | 0.00 | 0 | 0.000 | 0.0000 | 0.0000 | 0.0000 | 0.0000 |

In the search for unique features present in our individual, we followed the variation to genomic regions that have comparatively different SNP alleles relative to a representative sampling. The genomic regions are different from all of the comparison populations included in the study or from subsets. As it turns out, the SNPs directed the focus to regions of concentrated density and these regions might have something to tell. Lists of the genes that contain the distinctive elements that were found provide a further suggestion of how the natural process is working to provide advantage to our individual in the equally unique environment that surrounds him. As a better understanding of the prime function of these genes and gene interactions may emerge and as additional like individuals are characterized, the connections between distinctive genomic features and distinctive health features should be achieved.

## ACKNOWLEDGMENT

We acknowledge the support of King Abdullah International Medical Research Center, Ministry of National Guard Health Affairs, Riyadh, Kingdom of Saudi Arabia.

## AUTHOR CONTRIBUTIONS

I.A. and M.B. have contributed equally and were involved in the analysis and study design, coordination with the collaborators, organizing the study plan and preparing the manuscript. A.R. performed much of the analysis and contributed to the manuscript. S.A. was consulted in the study design, coordinates with the collaboration and contributed to the manuscript. M.J. as consultant in the overall study design and contributed to the manuscript. S.G. was involved in the wet laboratory work. M.B. was involved in the analysis and validation; organizing and supervising the over all work.

## COMPETING INTERESTS AND PERMISSIONS

The authors declare no competing interests and permissions.

## FUNDING AND PRIVACY

This work was funded by King Abdullah International Medical Research Center, Ministry of National Guard Health Affairs, Riyadh, Kingdom of Saudi Arabia.

## SUPPLEMENTAL MATERIALS

**Supplemental Table 1:** Affymatrix Compared to Sequencing for SNP detection

| chr | agreement | total_Affy SNPs | sequence SNPs | % Affy SNPs variable |
| --- | --- | --- | --- | --- |
| 1 | 0.916 | 27,777 | 226,896 | 38.9 |
| 2 | 0.923 | 29,326 | 239,542 | 39.7 |
| 3 | 0.926 | 24,167 | 198,277 | 39.8 |
| 4 | 0.916 | 23,141 | 219,531 | 41.3 |
| 5 | 0.918 | 22,408 | 180,453 | 39.7 |
| 6 | 0.916 | 21,613 | 186,211 | 38.4 |
| 7 | 0.914 | 18,560 | 164,975 | 39.4 |
| 8 | 0.922 | 19,595 | 157,322 | 40.3 |
| 9 | 0.930 | 15,899 | 123,218 | 38.3 |
| 10 | 0.921 | 18,845 | 151,480 | 39.1 |
| 11 | 0.921 | 17,826 | 147,017 | 40.0 |
| 12 | 0.924 | 16,590 | 134,261 | 39.0 |
| 13 | 0.934 | 13,904 | 111,491 | 40.5 |
| 14 | 0.921 | 11,372 | 92,810 | 29.9 |
| 15 | 0.932 | 10,814 | 82,667 | 41.4 |
| 16 | 0.918 | 11,834 | 94,347 | 41.1 |
| 17 | 0.935 | 8,226 | 75,332 | 39.8 |
| 18 | 0.926 | 10,873 | 86,779 | 41.0 |
| 19 | 0.918 | 4,709 | 60,106 | 39.5 |
| 20 | 0.918 | 8,393 | 63,417 | 36.7 |
| 21 | 0.912 | 4,896 | 46,565 | 38.9 |
| 22 | 0.904 | 4,155 | 38,003 | 36.0 |
| X | 0.991 | 9,388 | 64,308 | 25.1 |
| Y | 0.985 | 65 | 13,690 | 15 |
| M | NA | 0 | 31 | NA |
|  | **0.927** | average |  |  |

**Supplemental Table 2:** SNP and population allele sampling size

| chr | SNPs | AA<br>change | oSNPs | oAA<br>change | Ceu range;<br>avg | Chb range;<br>avg | Jpt range;<br>avg | Yri range;<br>avg |
| --- | --- | --- | --- | --- | --- | --- | --- | --- |
| 1 | 226,896 | 1,375 | 95,118 | 541 | 72-165; 160 | 30-48; 80 | 32-86; 81 | 72-67; 163 |
| 2 | 239,542 | 965 | 106,184 | 405 | 72-165; 124 | 30-48; 62 | 32-86; 62 | 72-167; 125 |
| 3 | 198,277 | 517 | 83,666 | 339 | 72-165; 126 | 29-48; 63 | 30-86; 63 | 72-167; 127 |
| 4 | 219,513 | 392 | 84,230 | 262 | 72-165; 123 | 31-48; 61 | 31-86; 62 | 72-167; 124 |
| 5 | 180,453 | 351 | 79,603 | 271 | 72-165; 125 | 31-48; 63 | 29-86; 63 | 72-167; 126 |
| 6 | 186,211 | 521 | 82,166 | 349 | 72-165; 124 | 29-48; 62 | 30-86; 62 | 72-167; 124 |
| 7 | 164,975 | 457 | 66,879 | 256 | 72-165; 125 | 29-48; 62 | 29-86; 63 | 72-167; 125 |
| 8 | 157,322 | 268 | 71,302 | 172 | 72-165; 124 | 31-48; 62 | 29-86; 62 | 72-167; 124 |
| 9 | 123,218 | 310 | 55,858 | 190 | 72-165; 125 | 30-48; 62 | 29-86; 63 | 72-167; 126 |
| 10 | 151,480 | 382 | 65,289 | 244 | 72-165; 124 | 30-48; 62 | 30-86; 62 | 72-167; 126 |
| 11 | 147,017 | 642 | 63,284 | 428 | 72-165; 124 | 32-48; 63 | 30-86; 63 | 72-167; 126 |
| 12 | 134,261 | 473 | 56,994 | 311 | 72-165; 126 | 30-48; 63 | 29-86; 63 | 72-167; 127 |
| 13 | 111,491 | 151 | 51,689 | 104 | 72-165; 121 | 30-48; 60 | 30-86; 61 | 72-167; 124 |
| 14 | 92,810 | 362 | 41,135 | 204 | 72-165; 124 | 32-48; 62 | 27-86; 63 | 72-167; 126 |
| 15 | 82,667 | 300 | 36,887 | 212 | 72-165; 126 | 30-48; 63 | 32-86; 64 | 72-167; 128 |
| 16 | 94,347 | 331 | 35,910 | 197 | 72-165; 128 | 30-48; 64 | 30-86; 65 | 72-167; 130 |
| 17 | 75,332 | 546 | 28,902 | 322 | 72-165; 130 | 30-48; 65 | 31-86; 65 | 72-167; 131 |
| 18 | 86,779 | 155 | 39,313 | 109 | 72-165; 123 | 31-48; 62 | 30-86; 62 | 72-167; 125 |
| 19 | 60,106 | 688 | 17,805 | 432 | 72-165; 133 | 32-48; 67 | 29-86; 67 | 72-167; 133 |
| 20 | 63,417 | 185 | 27,490 | 134 | 72-165; 128 | 32-48; 64 | 30-86; 65 | 72-167; 129 |
| 21 | 46,565 | 126 | 16,774 | 80 | 72-165; 127 | 31-48; 63 | 31-86; 64 | 72-167; 127 |
| 22 | 38,003 | 218 | 14,489 | 141 | 72-165; 130 | 31-48; 65 | 32-86; 65 | 72-167; 130 |
| X | 64,308 | 136 | 24,458 | 82 | 72-165; 129 | 31-48; 64 | 32-86; 64 | 72-167; 131 |
| Y | 13,690 | 1 | 69 | 0 | 40-45; 45 | 18-23; 23 | 20-23; 23 | 52-53; 53 |
| M | 31 | 7 | 0 | 0 | 83-165; 107 | 40-84; 52 | 38-86; 52 | 72-167; 118 |

Supplemental Figure 1: A decision formula was developed to call the observed SNP different from the corresponding SNP within a comparison population. The variables are calculated directly from the frequency data for each SNP. The decision is generated separately for homozygous or heterozygous alleles that are observed in the sample. Output is a decision of same or different.

